# Multiple invasions, *Wolbachia* and human-aided transport drive the genetic variability of *Aedes albopictus* in the Iberian Peninsula

**DOI:** 10.1101/2022.07.11.499514

**Authors:** Federica Lucati, Sarah Delacour, John R.B. Palmer, Jenny Caner, Aitana Oltra, Claudia Paredes-Esquivel, Simone Mariani, Santi Escartin, David Roiz, Francisco Collantes, Mikel Bengoa, Tomàs Montalvo, Juan Antonio Delgado, Roger Eritja, Javier Lucientes, Andreu Albó Timor, Frederic Bartumeus, Marc Ventura

## Abstract

The Asian tiger mosquito, *Aedes albopictus*, is a highly invasive species that has been spreading rapidly throughout tropical and temperate regions worldwide since the late 1970s. On the Iberian Peninsula, it was first recorded in 2004 near Barcelona. Since then, the species has spread along the Mediterranean coast and is now colonising the Peninsula’s inner territories. As with other species of the genus *Aedes*, the spread of the tiger mosquito has been linked to global shipping routes and road networks. In particular, the transport of adult mosquitoes by car has been shown to augment its natural spreading capacity by orders of magnitude. Much remains unknown, however, about the impact of human-mediated dispersal on the genetic variability of this species. This study aimed to ascertain the factors that have contributed to the spread and the current genetic variation of *Ae. albopictus* across the Iberian Peninsula, through complex dispersal mechanisms and endosymbiosis with bacteria like *Wolbachia*. We do so through population genetic analysis of mitochondrial (COI) and nuclear (ITS2) DNA sequences. Overall, both COI and ITS2 markers showed a lack of genetic structure among sampled regions and the presence of worldwide dominant haplotypes, suggesting a pattern of multiple introductions from abroad. We found extremely low levels of variation in COI compared to ITS2, and this lack of mitochondrial polymorphism is likely explained by high *Wolbachia* prevalence (79%). Multilevel models revealed that greater mosquito fluxes (estimated from commuting patterns and tiger mosquito population distribution) and spatial proximity between sampling sites were associated with lower ITS2 genetic distance, suggesting that rapid short- and medium-distance dispersal is facilitated by humans through vehicular traffic. This study highlights the significant role of human transportation in shaping the genetic attributes of *Ae. albopictus* and promoting regional gene flow, and underscores the need for a territorially integrated surveillance across scales of this disease-carrying mosquito.

**Author Summary:** The tiger mosquito, *Aedes albopictus*, is one of the most invasive species in the world. Native to the tropical forests of Southeast Asia, over the past 30 years it has rapidly spread throughout tropical and temperate regions of the world. Its dramatic expansion has resulted in public health concerns as a consequence of its vector competence for at least 16 viruses. Previous studies showed that *Ae. albopictus* spread has been facilitated by human-mediated transportation, but much remains unknown about how this has affected its genetic attributes. Here we examined the factors that contributed to shaping the current genetic constitution of *Ae. albopictus* in the Iberian Peninsula, where the species was first found in 2004, by combining population genetics and Bayesian modelling. We found that both mitochondrial and nuclear DNA markers showed a lack of genetic structure and the presence of worldwide dominant haplotypes, suggesting regular introductions from abroad. Mitochondrial DNA showed little genetic diversity compared to nuclear DNA, likely explained by infection with maternally transmitted bacteria of the genus *Wolbachia*. Our models indicated that human transportation plays a role in shaping *Ae. albopictus* nuclear genetic structure by means of passive dispersal of adult tiger mosquitoes through the road network.

## Introduction

The Asian tiger mosquito, *Aedes* (*Stegomyia*) *albopictus* (Skuse 1894) is a highly invasive species that originated in the tropical forests of Southeast Asia [1]. However, in the late 1970s it started a dramatic expansion throughout tropical and temperate regions of the world and it is now present in the five populated continents [2]. This species is an ecological generalist capable of rapid evolution and, with the aid of man, speedy colonization of new habitats [1]. While in its native range *Ae. albopictus* inhabits forested areas, breeding in natural sites such as tree holes, bromeliads and bamboo stumps, it has now adapted to breed also in artificial man-made water containers from urban and suburban human settlements [3]. This species has an opportunistic feeding behaviour with a strong preference for mammals, especially humans [4, 5]. Furthermore, in temperate regions, its eggs can survive the cold winters by entering diapause [3]. This ecological adaptability has important implications for the epidemiology of several mosquito-borne diseases since the tiger mosquito is a competent laboratory vector of at least 16 viruses [5, 6]. While *Ae. aegypti* is considered as the principal vector of dengue and Zika, *Ae. albopictus* is a less efficient epidemic vector (with local exceptions in some cases), having developed an enhanced transmission for chikungunya facilitated by genetic adaptation (E1-A226V substitution) of the ECSA strain [6–9]. *Ae. albopictus* has a role on sporadic autochthonous disease transmission of arboviruses (as chikungunya, dengue and Zika) in Europe, and therefore its surveillance and control are considered a regional priority [10]. Invasive alien species have historically been spread by humans. However, advances in transportation logistics resulting in higher air traffic and sea-born trading are driving a more rapid dispersal of non-indigenous species in the world, including many vectors of human diseases [11]. *Ae. albopictus* is one of top invasive alien species in the world, being considered, together with *Ae. aegypti*, the most costly invasive species [12, 13]. Its expansion in Europe coincides with the wave of introductions of invasive species in the continent, which started almost 40 years ago as a result of globalization [14]. Considering its limited flight range (less than 200 meters/day) [15], a crucial element for its success is the longevity of its desiccation-resistant eggs [16], which can be passively transported by humans through commercial shipping of used tires and aquatic plants [3] and ground vehicles [10, 17]. In Spain, *Ae. albopictus* was first detected in Sant Cugat del Vallès, Catalonia, in 2004 [18] as a result of the nuisances associated with its aggressive anthropophilic behaviour [19]. Almost twenty years later, this species is well established in the Mediterranean coast of Spain and it is now colonizing inner territories of the Iberian Peninsula [17, 20].

A biological invasion is a three-step process, which involves: initial dispersal, establishment of self-sustaining populations and spread to neighbouring habitats [21, 22]. It is, however, during the initial dispersal when proactive management efforts can be more cost-effective for preventing the establishment of invasive species [22, 23]. For instance, successful surveillance in New Zealand in the 1990s intercepted the entrance of *Ae. albopictus* [24], which has not been reported so far in this country. On the other hand, once this has become established, it is highly difficult to eradicate [16]. In this respect, studying *Ae. albopictus* population genetic structure and identifying its dispersal routes, its main drivers and scales, is crucial to understand the tiger mosquito’s spread and design integrated surveillance and preparedness strategies [25, 26]. Previous population genetic studies pointed to a worldwide chaotic dispersion pattern in *Ae. albopictus*, with Europe harbouring several distinct genotypes that have been linked to multiple independent introductions [e.g. 25, 27, 28]. Furthermore, in Europe human transportation networks have been shown to have facilitated the spread of this mosquito and conditioned its genetic and demographic patterns [29–31].

As different marker types can tell different stories [32, 33], it is crucial to analyse both mitochondrial (mtDNA) and nuclear (nDNA) DNA data to understand what major factors contributed to shaping the genetic attributes of invasive populations. In this regard, the presence of maternally-transmitted endosymbiotic bacteria may be relevant, as they can distort mtDNA phylogenies and reduce mtDNA diversity, a process which may have little to no effects on nDNA variation [34]. If a maternally-inherited symbiont confers a selective advantage to its hosts, the mtDNA variants originally associated with the symbiont can rapidly spread through the host population and go to fixation, thus resulting in a great increase in the frequency of a single or few mtDNA haplotypes [35]. Such symbionts are widespread in many arthropod species, being *Wolbachia* the most common maternally-inherited symbiont on the planet [36, 37]. For this reason, incorporation of nDNA data is essential to corroborate results derived from mtDNA.

It is now widely accepted that a progress in the study of the ecology of *Ae. albopictus* requires an interdisciplinary approach [11, 38]. For instance, population genetics and multilevel modelling analyses can be crucial to shed light on *Ae. albopictus* patterns of dispersal. Besides, it has been shown that genetic diversity in this species has been related to a higher adaptive potential [29] and a diversification of its interactions with the pathogens it carries [38]. Understanding movement patterns in its populations in endemic areas is crucial to disentangle the dynamics of disease transmission between vectors and humans [15]. In this study we assess geographic patterns of mtDNA and nDNA variation in *Ae. albopictus* and use this information to investigate the species’ dispersal routes across Spain. We also compare features of mtDNA and nDNA variation and assess whether they are incongruent, and test for *Wolbachia* presence to ascertain whether mtDNA diversity is affected by this maternally-transmitted parasite.

## Materials and Methods

### Sample collection and DNA extraction

*Ae. albopictus* samples were collected during the period 2011-2015 at 140 locations encompassing most of the current species distribution in Spain (13 provinces mostly located along the Mediterranean coast; Fig 1, Table 1). Additional samples were collected from two areas of France (Nice N 43° 38’ 32’’ E 7° 5’ 27’’ and Montpellier N 43° 40’ 39’’ E 4° 2’ 35’’) and two regions of Greece (the area of Pylaia, in Salonica N 40° 27’ 22’’ E 23° 13’ 20’’ and Sykia N 40° 2’ 19’’ E 23° 56’ 22’’), which were used as complementary data from these Mediterranean areas (Table 1). Sample collection was performed using different methods according to the life stage (adults, larvae or eggs). Adult tiger mosquitoes were collected by citizens who participated in the *Mosquito Alert* programme (see below). Mosquito larvae and eggs were either collected from road drains or from standard ovitraps, which consisted of a dark plastic container filled with water and a wooden board as oviposition support. Ovitraps were positioned in shaded sites in petrol stations, cemeteries, private properties and public parks. They were inspected weekly or biweekly, wooden boards replaced and the water replenished. Eggs were hatched in the laboratory of the Centre d’Estudis Avançats de Blanes (CEAB-CSIC, Spain), reared to the fourth-larval instar or adult stage and then identified following Schaffner et al. [39]. All samples were stored in absolute ethanol and kept at -20°C until genetic analyses.

**Fig 1.**
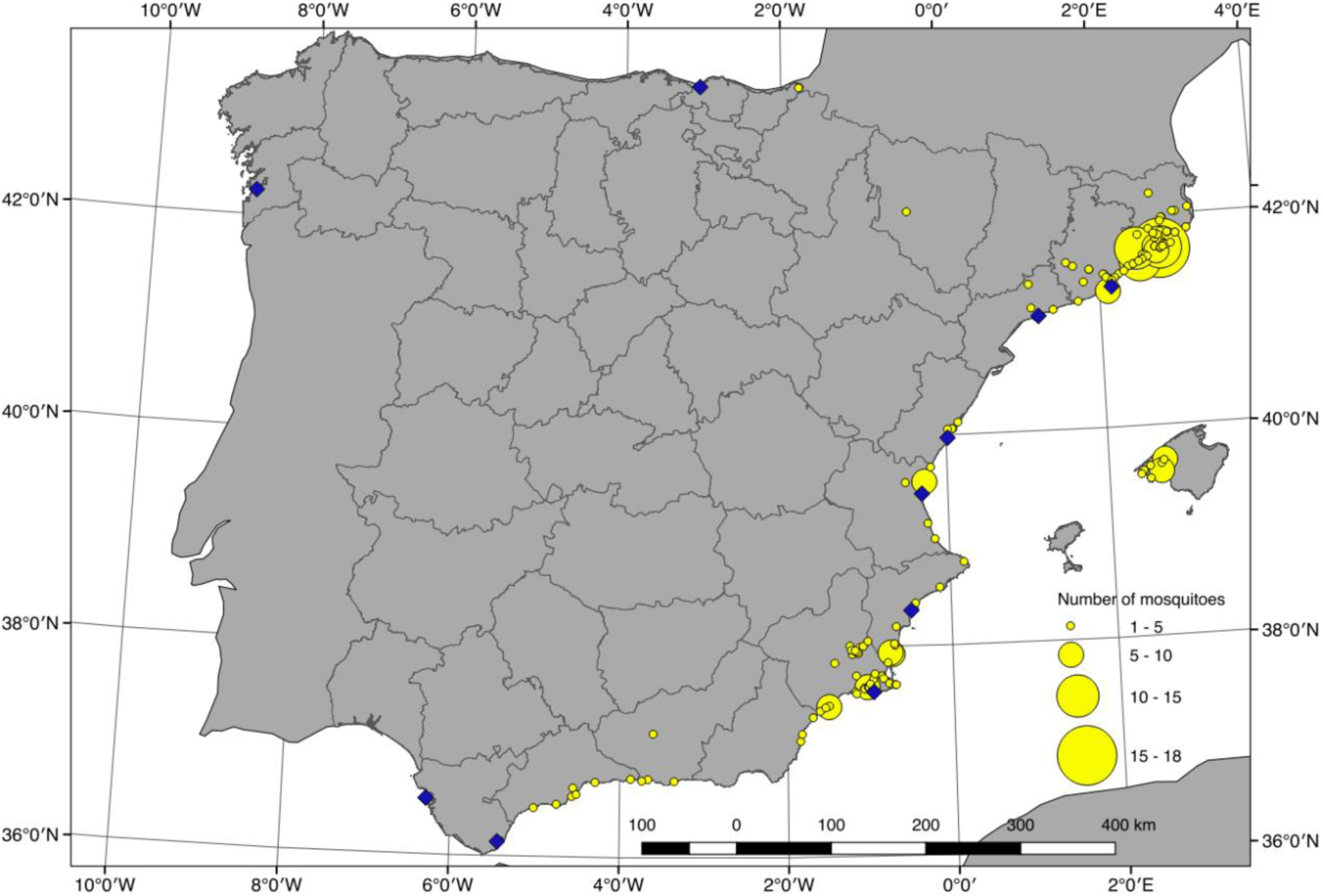
Localities sampled for genetic analysis. Yellow circles are centered on sample locations with radius proportional to number of mosquitoes sampled at each location. Blue diamonds indicate locations of major commercial ports in Spain (ports with over 50,000 TEU -Twenty-Foot Equivalent Unit-, based on AAPA -American Association of Port Authorities- data from 2013). Grey lines indicate provinces of Spain. Land boundaries from Natural Earth. Province boundaries from GADM.

**Table 1.** Basic genetic statistics of Ae. albopictus sampled provinces for mtDNA (COI) and nuclear DNA (ITS2).

| Province/Country | Year of first detection | Sampling period | N<sub>COI</sub> | S<sub>COI</sub> | HD<sub>COI</sub><br>(mean ± SE) | π<sub>COI</sub><br>(mean ± SE) | N<sub>ITS2</sub> | S<sub>ITS2</sub> | HD<sub>ITS2</sub><br>(mean ± SE) | π<sub>ITS2</sub><br>(mean ± SE) |
| --- | --- | --- | --- | --- | --- | --- | --- | --- | --- | --- |
| Barcelona | 2004 | 2012-2015 | 104 | 10 | 0.262 ± 0.055 | 0.0008 ± 0.0003 | 104 | 32 | 0.693 ± 0.050 | 0.005 ± 0.001 |
| Girona | 2008 | 2011-2015 | 80 | 5 | 0.233 ± 0.062 | 0.0005 ± 0.0003 | 84 | 54 | 0.597 ± 0.059 | 0.006 ± 0.002 |
| Tarragona | 2005 | 2012-2015 | 7 | 1 | 0.286 ± 0.196 | 0.0006 ± 0.0005 | 4 | 2 | 0.833 ± 0.222 | 0.004 ± 0.002 |
| Mallorca | 2012 | 2014-2015 | 40 | 2 | 0.145 ± 0.074 | 0.0003 ± 0.0002 | 33 | 35 | 0.818 ± 0.067 | 0.012 ± 0.003 |
| Huesca | 2015 | 2015 | 3 | 0 | 0 | 0 | 5 | 10 | 1.000 ± 0.126 | 0.014 ± 0.005 |
| Castelló | 2010 | 2011-2015 | 7 | 1 | 0.286 ± 0.196 | 0.0006 ± 0.0005 | 7 | 5 | 0.524 ± 0.209 | 0.007 ± 0.003 |
| València | 2013 | 2011-2015 | 20 | 0 | 0 | 0 | 18 | 17 | 0.490 ± 0.142 | 0.006 ± 0.002 |
| Alacant | 2006 | 2011-2015 | 16 | 4 | 0.350 ± 0.148 | 0.0014 ± 0.0007 | 34 | 17 | 0.729 ± 0.078 | 0.005 ± 0.002 |
| Murcia | 2011 | 2012-2015 | 110 | 2 | 0.240 ± 0.049 | 0.0005 ± 0.0004 | 111 | 15 | 0.658 ± 0.031 | 0.004 ± 0.002 |
| Málaga | 2014 | 2014-2015 | 20 | 1 | 0.190 ± 0.108 | 0.0004 ± 0.0004 | 10 | 26 | 0.978 ± 0.059 | 0.020 ± 0.005 |
| Granada | 2014 | 2014-2015 | 4 | 0 | 0 | 0 | 6 | 14 | 0.800 ± 0.172 | 0.022 ± 0.007 |
| Almería | 2014 | 2014-2015 | 7 | 0 | 0 | 0 | 6 | 4 | 0.600 ± 0.215 | 0.005 ± 0.002 |
| Gipuzkoa | 2014 | 2014-2015 | 6 | 0 | 0 | 0 | 2 | 19 | 1.000 ± 0.500 | 0.064 ± 0.016 |
| France | - | 2014-2015 | 40 | 0 | 0 | 0 | 30 | 22 | 0.867 ± 0.043 | 0.012 ± 0.003 |
| Greece | - | 2013-2014 | 7 | 0 | 0 | 0 | 16 | 13 | 0.617 ± 0.135 | 0.006 ± 0.002 |

N, sample size; S, number of polymorphic sites; HD, haplotype diversity; π, nucleotide diversity.

DNA was extracted from the whole bodies of mosquitoes (adults or larvae) in a final volume of 250 µL using the HotShot protocol [40]. Briefly, samples were incubated with 125 ml of Alkaline lysis buffer for 30 min at 96 °C, which were afterwards neutralized with 125 ml of neutralizing solution (Tris-HCl 40 mM, pH 5.0), and subsequently stored at -20°C.

### *Ae. albopictus* nuclear and mitochondrial gene sequencing

We amplified two gene regions, including one nuclear ribosomal gene (the second internal transcribed spacer of ribosomal DNA - ITS2) and one mitochondrial fragment (the cytochrome *c* oxidase gene subunit 1 - COI). The following primers were used for amplification and sequencing: for ITS2, primers ITS-CP-P1A (5’- GTGGATCCTGTGAACTGCAGGACACATG-3’) and ITS-CP-P1B (5’- GTGTCGACATGCTTAAATTTAGGGGGTA-3’) [41], and for COI, primers LCOI490 (5′-GGTCAACAAATCATAAAGATATTGG-3′) and HCO2198 (5′-TAAACTTCAGGGTGACCAAAAAATCA-3′) [42] or the degenerated primers ZplankF1_M13 (5’-TGTAAAACGACGGCCAGTTCTASWAATCATAARGATATTGG-3’) and ZplankR1_M13 (5’-CAGGAAACAGCTATGACTTCAGGRTGRCCRAARAATCA-3’) [43] that had the M13 primers attached following the suggestion of Ivanova et al. [44]. Amplifications were done with a total reaction volume of 25 µL, containing 1× PCR buffer (Silverstar, Eurogentec), 1.5 mm MgCl2, 200 µm of each dNTP, 0.2 µM of each primer, 7 µL of template DNA for ITS2 and 2 µL for COI, 1 U Taq polymerase and UV light-sterilized mQ-H2O. PCR amplification for ITS2 primers involved a denaturing step of 1 min at 94°C, five cycles of 40 s at 94°C, 2 min at 37°C and 60 s at 72°C, followed by 35 cycles of 40 s at 94°C, 40 s at 51°C and 60 s at 72°C and a final elongation step of 6 min at 72°C. For COI Folmer et al. [42] primers, PCR amplification involved a denaturing step of 5 min at 95°C, five cycles of 60 s at 95°C, 90 s at 45°C and 45 s at 72°C, followed by 30 cycles of 45 s at 95°C, 45 s at 50°C and 45 s at 72°C and a final elongation step of 7 min at 72°C, while for COI-degenerated primers amplification conditions consisted of 35 cycles of 1 min at 95°C, 2 min at 37°C and 1 min at 72°C. PCR products were purified and sequenced on a ABI 3730XL capillary sequencer by a third party (Macrogen, Seoul, South Korea). In the case of ITS2, 22.7% (138) of the individuals could not be considered for analysis due to intragenomic heterogeneity (simultaneous presence of two or more haplotypes in the same individual). Resulting sequences were aligned with other relevant *Ae. albopictus* sequences from each gene (retrieved from GenBank) using the ClustalW algorithm in MEGA 7 [45].

### Wolbachia screening

To test whether mtDNA variability could be affected by the presence of endosymbiotic bacteria of the genus *Wolbachia*, we analysed 62 randomly selected samples (13.2% of the overall samples). Individuals were selected aiming to cover all sampling years and all studied provinces. Two molecular markers were used for detecting *Wolbachia* infection, namely *wsp* and 16S rDNA. Primers used for amplification and sequencing were: for the *wsp* marker, primers 81F (5’- TGGTCCAATAAGTGATGAAGA-3’) and 691R (5’- AAAAATTAAACGCTACTCCA-3’) [46, 47], while for 16S, primers 16SF (5’- CGGGGGAAAAATTTATTGCT-3’) and 16SR (5’-AGCTGTAATACAGAAAGTAAA-3’) [48, 49]. Amplification conditions followed Wiwatanaratanabutr [50] in the case of *wsp*, and Heddi et al. [49] in the case of 16S. PCR products were visualized on a 1% agarose gel. To validate the results, two PCR replicates were run for each sample (following Carvajal et al. [51]). A third replicate was run for samples that showed incongruent results based on the two prior replicates. *Wolbachia* infection was confirmed by two successful amplifications of both molecular markers. In light of the high infection rate detected in the selected samples (see Results), we decided to exclude COI from population structure and multilevel modelling analyses (see below), as mtDNA diversity was likely affected by the presence of *Wolbachia*.

### Genetic diversity and structure

Genetic diversity indices including number of haplotypes (H), number of polymorphic sites (S), haplotype diversity (HD) and nucleotide diversity (π) for mtDNA and nuclear DNA were estimated for the whole dataset and for each analysed province using DNASP 6.11.01 [52]. To estimate gene genealogies, we used HAPLOVIEWER, which turns trees built from traditional phylogenetic methods into haplotype genealogies [53]. We estimated the phylogeny using a maximum-likelihood approach as implemented in RAxML 7.7.1 [54], with a GTRCAT model rate of heterogeneity and no invariant sites for COI, and a gamma model rate of heterogeneity and invariant sites (GTR+G+I) for ITS2. The most appropriate model of nucleotide evolution was selected using jModelTest 2.1.3 [55] under the Akaike information criterion (AIC). Input data were COI or ITS2 sequences from each individual, subsequently collapsed into haplotypes. The best tree was selected for network construction in HAPLOVIEWER.

Population structure of nuclear DNA was characterized using classical multidimensional scaling (MDS), also known as principal coordinates analysis [56]. Pairwise distances between ITS2 sequences were calculated in MEGA by estimation of evolutionary divergence over sequence pairs, using the Kimura 2-parameter substitution model. MDS analysis was then performed with the pairwise genetic distance matrix, using the cmdscale function of the stats package in R 4.1.2 [57].

### Construction of covariates

Our analysis of the drivers of genetic distance focused on four constructed covariates: spatial distance, spatial proximity, temporal distance, and potential tiger mosquito flux. We constructed the spatial distance variable as the geodesic distance between sample locations (inter-point distance) in meters, calculated using the Vincenty method on the WGS84 ellipsoid as implemented in the geosphere package for R [58]. We constructed spatial proximity as the negative exponential of spatial distance in km, represented as e^-d^, where e is Euler’s constant and d is spatial distance in km. This approach gives high values for spatial proximity (0.82-1) within the 200m buffer often taken as the tiger mosquito’s maximum flying distance, with values then quickly dropping and nearing zero by 5 km. The combination of these spatial distance and spatial proximity variables together is intended to capture the effects of inter-point distance at different scales. We constructed the temporal distance variable as the absolute number of years elapsed between the sampling times when each member of the sample pair was captured, rounded up to the nearest year (another way of thinking about this is as the number of mosquito seasons between each capture time). We constructed potential tiger mosquito flux as the potential daily bidirectional gross number of *Ae. albopictus* moving between each pair of municipalities. We estimated this by combining municipality-level *Ae. albopictus* risk estimates from the *Mosquito Alert* citizen science platform [59] with commuter flow estimates drawn from the Spanish Labour Force Survey (LFS) in a manner similar to that described in Eritja et al. [30]. The *Ae. albopictus* risk estimates were made with a Bayesian multilevel logistic regression of *Ae. albopictus* presence in Spain measured as expert-validated *Mosquito Alert* reports of adult *Ae. albopictus* at given points in time and space from 2014 through 2021. We combined these reports of presence with pseudoabsences created by placing points randomly within the same geographic and temporal bounds. The space-time region within these bounds was divided into sampling-cell-days, using the 0.025 degree latitude by 0.025 degree longitude sampling cells within which the *Mosquito Alert* app collects anonymous background tracks and estimates sampling effort based on the number of citizen scientists in each cell and the amount of time elapsed since each downloaded the app (since participant motivation has been observed to drop over time) [59]. The number of pseudoabsences placed in each sampling-cell-day was proportional to the *Mosquito Alert* sampling effort. The model includes random intercepts for municipalities and province-years to capture spatio-temporal variation in the observed *Ae. albopictus* distribution. It includes random intercepts for landcover, constructed from the 2018 Corine Landcover dataset, and it includes a random slope at the province level for the Mosquito Weather Index, developed by the LIFE CONOPS project [60] and constructed using the mean monthly temperature, humidity, and wind velocity variables from the ERA5 dataset [61]. The model also includes an offset for sampling effort and random intercepts at the sampling cell level to control for variation and anomalies in sampling behaviour by the citizen scientists beyond what is captured by the placement of pseudoabsences.

The *Ae. albopictus* risk estimates were combined with the estimated daily number of people commuting between each pair of municipalities in either direction. These commuting estimates were made using data from the LFS, a continuous survey of the Spanish population living in family dwellings, which is administered four times per year using a two-stage stratified random sample with approximately 160,000 respondents per wave [62]. The survey includes questions about each respondent’s home province and the province in which they work, which can be used to estimate commuter fluxes between provinces [30, 63]. Inter-province commuter fluxes were estimated on a quarterly basis directly from the LFS micro-data using the survey weights to make population level estimates, and calculating mean fluxes per year. These estimates were then down-scaled to the municipality level using administrative data on places of domicile. Inter-province fluxes were distributed across municipalities in proportion to the relative residential populations in each municipality, based on the assumption that inter-province commuters’ homes are distributed across municipalities in their home province similarly to the rest of the population’s homes in that province, and that inter-province commuters’ places of work are distributed across municipalities in their work province similarly to the rest of the population’s homes in that province [30]. The inter-municipality fluxes are shown in S1 Fig.

### Modelling the influence of spatial proximity, mosquito flux, and temporal distance on genetic distance

We explored drivers of nuclear genetic structure by carrying out simple Mantel tests of correlations [64] between ITS2 genetic distance and potential tiger mosquito flux, spatial distance, spatial proximity, and temporal distance. We implemented all tests in the ecodist package for R [65], using 1,000 permutations with bootstrap confidence limits estimated using 500 iterations.

We then used Bayesian multiple membership multilevel regressions to model ITS2 pairwise genetic distances between sampled mosquito pairs as a function of potential tiger mosquito flux between sampling sites, spatial distance and spatial proximity between sampling sites, and temporal distance.

We relied on zero-inflated beta regressions [66], in which the distribution of the stochastic component is a mixture of a beta distribution and a degenerate distribution in 0. Following Figueroa-Zúñiga et al. [67] and Branscum et al. [68], we used the beta distribution for ITS2 genetic distance values in (0, 1) because it is a highly flexible distribution that is defined on that interval. As these authors have done, we parameterized the beta distribution in terms of a mean (*μ*) and a parameter (*ϕ*) that captures precision. The probability density function of a variable (*y*) in this parameterization is:

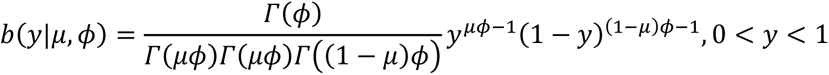

Since the beta distribution is undefined for values of zero, we used a zero-inflation component. As explained in Ospina and Ferrari [69], the probability density function of a variable *y* for this zero-inflated beta (zib) mixture is:

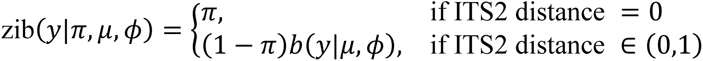

where 0<*π*<1 is the probability of observing an ITS2 distance of 0, and *μ* and *ϕ* are defined as above.

We treated the observed ITS2 genetic distances between each sample pair *i* as independent random variables *y1,…,yn* drawn from this zero inflated beta distribution such that *y*_*i*_ ∼ zib(*y*|*π*_*i*_, *μ*_*i*_, *ϕ*_*i*_). We used a logit link to model both *π* and *μ*, and a log link to model *ϕ*. We fit six models (M1-M6) with different combinations of mosquito flux, spatial proximity, temporal distance, and spatial distance as covariates. The models for *μ* are specified as following:

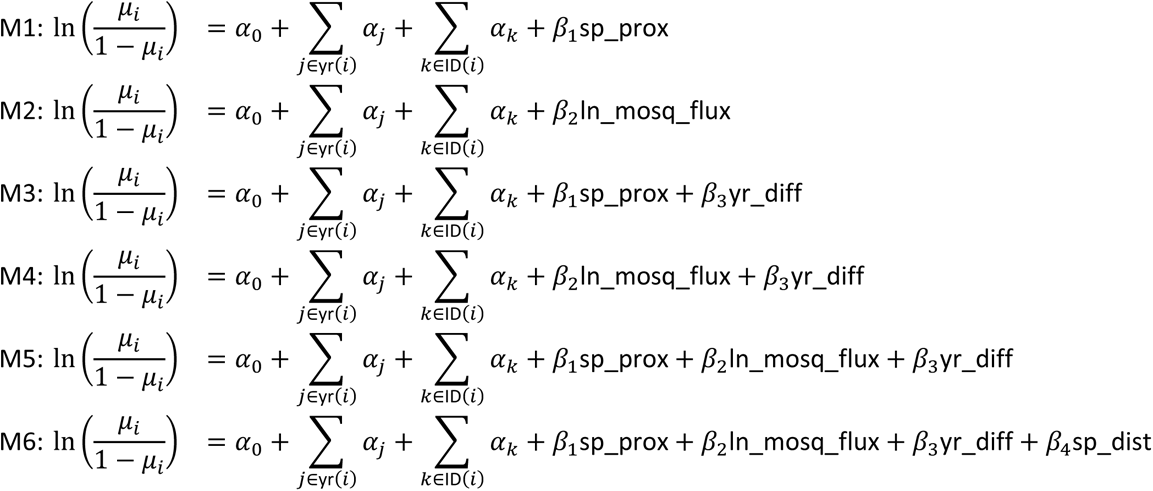

where *αj* represents a random intercept for each of the years in which the mosquitoes in pair *i* were sampled, and *αk* represents a random intercept for each of the sampled mosquitoes. This multiple membership approach allows us to account for the effects of sample year and sample itself, which could otherwise confound results given that each pair may be connected to other pairs by the sampling years and sampling that they share [70]. *β*_1_, *β*_2_, *β*_3_and *β*_4_are the slopes on the spatial proximity (sp_prox), log mosquito flux (ln_mosq_flux), temporal distance (yr_diff) and spatial distance (sp_dist) variables, and *α0* is the overall model intercept.

We explicitly model the *ϕ* parameter of our beta distributions as a function of the provinces in which each sample in the pair was taken. This allows us to explore how the precision of the beta distribution changes across geography. The choice of provinces as the areal units for this part of the model is based on these units representing patterns of human settlement and activity that should be of relevance to tiger mosquito spreading patterns, while also being large enough to avoid adding too many additional parameters to an already complicated model. The model for *ϕ* is the same in all six models:

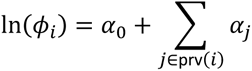

Finally, we modelled *π* (the probability of ITS2 pairwise distance being equal to zero) as:

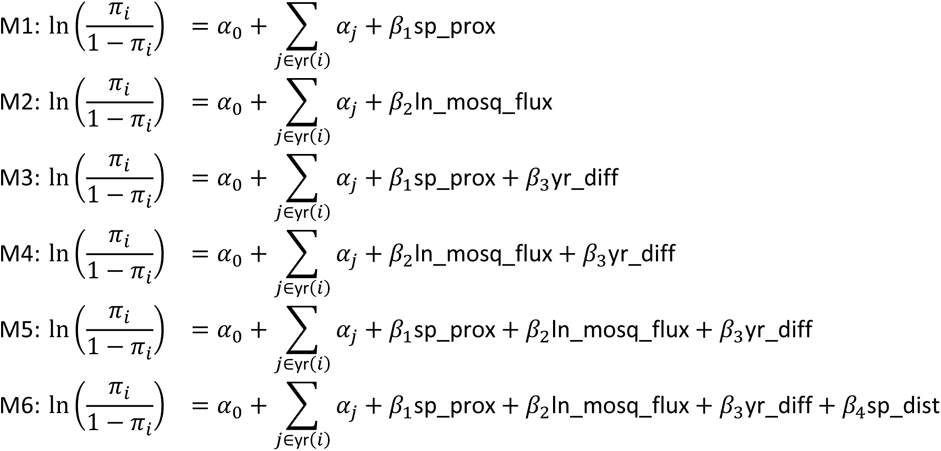

The systematic part of the model for *π* is almost identical to the systematic part of the model for *μ*. The only difference is the absence of the sample random intercepts, which are excluded here for computational reasons.

Models were fit using Hamiltonian Monte Carlo Markov (HMC) sampling implemented by Stan [71] using the brms package for R [72]. Computation was done on the CEAB-CSIC’s high-performance computational cluster, using 48 cores and within-chain parallelization implemented by the cmdstanr package for R [73]. All independent variables were centered and standardized before model fitting. We compared models using leave-one-out cross validation [LOO; 74, 75] and Bayesian R-squared [76]. Given the large size of the dataset, we carried out LOO using subsamples of the data in order to speed computation, as described in Magnusson et al. [74]. LOO was then estimated with the Pareto-smoothed importance sampling proposed by Vehtari et al. [75]. Expected log pointwise predictive density (ELPD) was used as a measure of predictive accuracy. This analysis was implemented using the loo [75, 77] and brms packages in R. As for the R-squared, we used the Bayesian version of R-squared proposed by Gelman et al. [76] to avoid the problem that the numerator may be larger than the denominator in Bayesian models. We implemented this using the brms package in R.

## Results

### Genetic variation and structure, and temporal trends of genetic diversity

The nuclear DNA alignment (ITS2) included 470 sequences of 286 bp (424 sequences from Spain + 30 from France + 16 from Greece), while for the mitochondrial DNA alignment (COI) we obtained 471 sequences of 513 bp (424 from Spain + 40 from France + 7 from Greece) (Table 1). Haplotype and nucleotide diversity estimates calculated at the province level ranged from 0.490 to 1 and from 0.004 to 0.064 for ITS2, respectively, and from 0 to 0.350 and from 0 to 0.0014 for COI (Table 1).

In the Iberian Peninsula, overall ITS2 genetic diversity was between three (haplotype diversity) and 10 (nucleotide diversity) times higher than for COI. Indeed, overall haplotype (HD) and nucleotide (π) diversities were 0.7084 ± 0.025 and 0.00566 ± 0.00048 for ITS2, respectively, and 0.2196 ± 0.027 and 0.00054 ± 0.00008 for COI. We found 78 haplotypes defined by 83 polymorphic sites (21 parsimony informative) for ITS2, whereas only 18 haplotypes defined by 20 polymorphic sites (5 parsimony informative) were detected in the case of COI (Fig 2).

**Fig 2.**
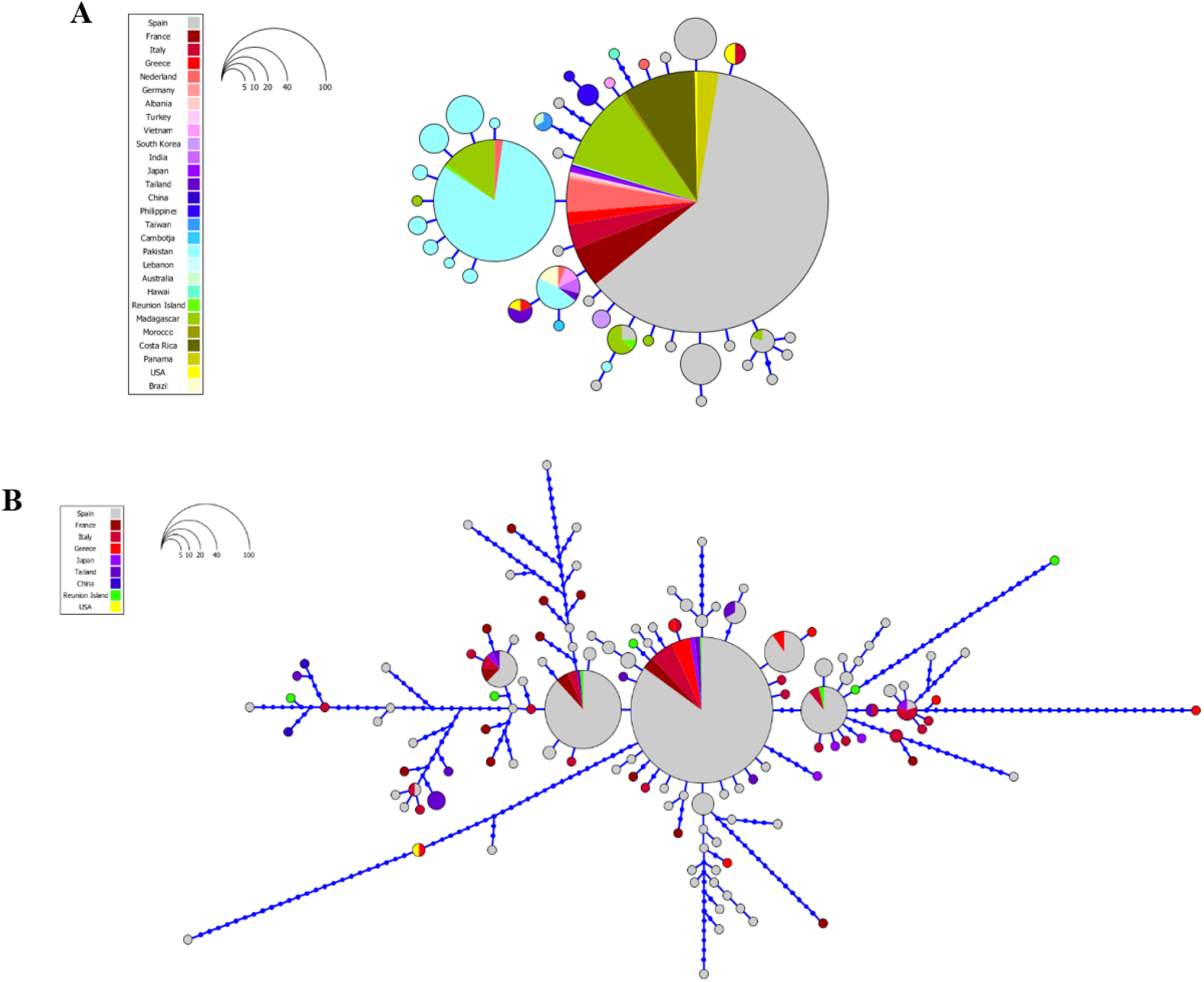
Haplotype networks of COI mtDNA sequences (A) and ITS2 nuclear DNA sequences (B) analysed in *Ae. albopictus*. Each circle represents a unique haplotype and the circle area is proportional to the number of sequences of a given haplotype. Blue dots correspond to inferred unsampled haplotypes.

No apparent association between haplotypes and geography was detected. Both ITS2 and COI haplotype networks showed the presence of worldwide dominant haplotypes and the absence of a clear genetic structure along the study area (Fig 2). Specifically, in the ITS2 network the most frequent and centrally placed haplotypes were observed in localities spanning the entire extension of the study area and were shared with other areas of the world. However, 89.7% of the haplotypes found were newly described (70/78), the majority being low-frequency haplotypes. In the case of COI, we obtained a much simpler star-like network defined by one main central haplotype connected to many low-frequency haplotypes with little genetic differentiation from the dominant sequence. Haplotypes were separated by no more than three nucleotide substitutions, although 83.3% of the haplotypes found were unique (15/18).

At the province level, we found a significant positive correlation between genetic diversity and colonisation time for COI, meaning that the provinces that were first colonised by *Ae. albopictus* bear higher mitochondrial genetic diversity (R^2^ = 0.497, p = 0.007 for haplotype diversity, R^2^ = 0.597, p = 0.002 for nucleotide diversity; Fig 3). However, a deeper look showed that COI’s variation in nucleotide diversity over time was almost non-existent (π range: 0-0.0014), likely indicating a statistically significant but biologically irrelevant pattern. On the contrary, we found a significant negative correlation between ITS2 nucleotide diversity and colonisation time (R^2^ = 0.376, p = 0.022), which displayed higher variation (π range: 0-0.022); the same relationship with haplotype diversity was not significant (Fig 3).

**Fig 3.**
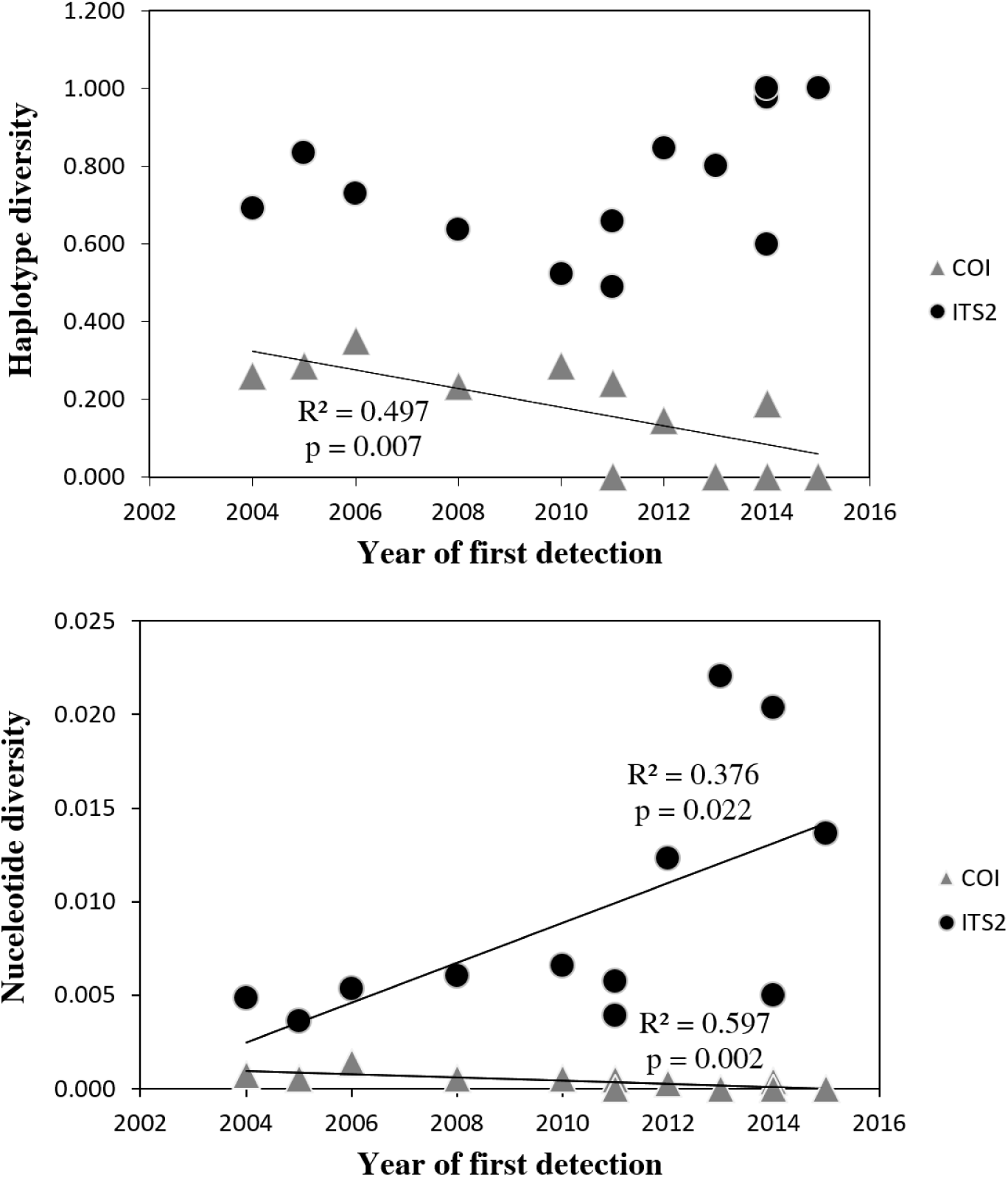
Relationship between genetic diversity and year of first detection in the analysed provinces. The mitochondrial COI fragment is indicated by grey triangles and the nuclear ITS2 gene by black circles. See Table 1 for province list.

In agreement with the haplotype network, MDS analysis revealed no apparent evidence of distinct genetic groups or geographic consistency within the nuclear dataset, although it did show a certain level of drift over time (Fig 4). Most of the samples were grouped into a unique cluster in the centre of the MDS, with the exception of a few samples collected in 2014 and 2015 at the geographic extremes of the study area that laid slightly outside the main cluster (i.e. samples from the southernmost provinces of Málaga, Granada and Murcia, from the northernmost provinces of Gipuzkoa and Girona, and from the Balearic Islands in the east). The two-dimensional MDS solution accounted for 30% of the variance in the data (goodness of fit = 0.302).

**Fig 4.**
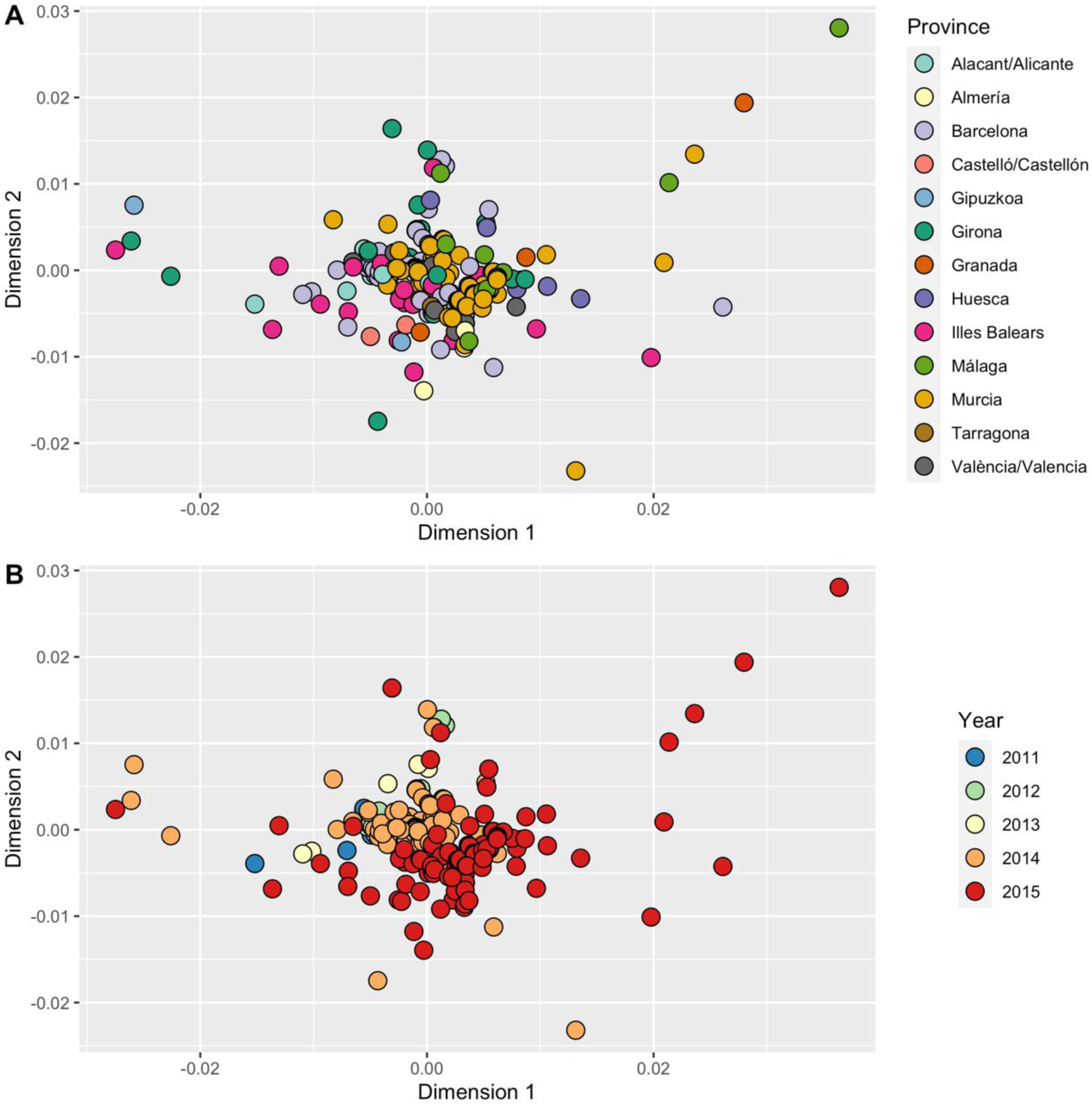
Multidimensional scaling plots of ITS2 genetic distances among *Ae. albopictus* samples, coloured by province (A) and year (B) of sample collection. Plots show the 2-dimensional solution using classical scaling. Goodness of fit = 0.302.

The Mantel test of correlation revealed a significant positive correlation between ITS2 pairwise genetic distances and spatial distance (r = 0.163, p = 0.001), and negative correlations with potential tiger mosquito flux (r = -0.022, p = 0.036) and spatial proximity (r = -0.042, p = 0.001). The Mantel test did not find evidence of correlation between ITS2 genetic distance and temporal distance.

### Wolbachia infection

From a total of 62 samples screened using the *wsp* and 16S markers, 49 (79%) from 44 localities tested positive for infection for both markers. Two additional individuals yielded positive amplifications for 16S only and were thus excluded in reporting *Wolbachia* prevalence.

### Modelling genetic distance from spatial distance, temporal distance and mosquito flux

Our multilevel models indicated that ITS2 genetic distance was better explained by the combination of potential tiger mosquito flux, spatial proximity, spatial distance and temporal distance than by any of these variables on their own or in smaller combinations. This is most clearly seen in our LOO comparison, in which the expected log pointwise predictive density for M6 is 138 points lower than the next best model, with a standard error of only 20 (S1 Table). It is also seen in the Bayesian R-squared comparison, in which M6 also has the highest value, although in this case the differences are small compared to the standard errors (complete model comparisons are presented in S2 Fig and S2 Table). The slope estimates for the main effects in the equation for the mean (*μ*) of the beta distribution are shown in Fig 5A and S2 Table, and should be read in conjunction with the slope estimates for the main effects in the equation for the probability of zeros (*π*), shown in Fig 5B and S2 Table. The coefficients on the spatial proximity and potential tiger mosquito flux variables are negative in all models of *μ* and positive in all models of *π*, even when both variables are included together (M5 and M6), indicating that greater spatial proximity and mosquito fluxes are associated with lower ITS2 genetic distance between sampled mosquitoes, with a higher probability of zero distances. In contrast, the coefficient on the temporal distance variable is positive in all models of *μ* and negative in all models of *π* (albeit with its posterior distribution overlapping zero in M5 and M6), indicating that greater passage of time between samples is associated with greater ITS2 genetic distance and with lower probability of zero distances (S3 and S4 Figs). In M6 we find that greater spatial distance is also associated with lower probabilities (*π*) of zero ITS2 genetic distance (Fig 5B). Although greater spatial distance is also associated with lower ITS2 genetic distance in the model of *μ* (Fig 5A), the combined effect of the two components is an overall positive relationship between spatial distance and ITS2 genetic distance (Fig 6A).

**Fig 5.**
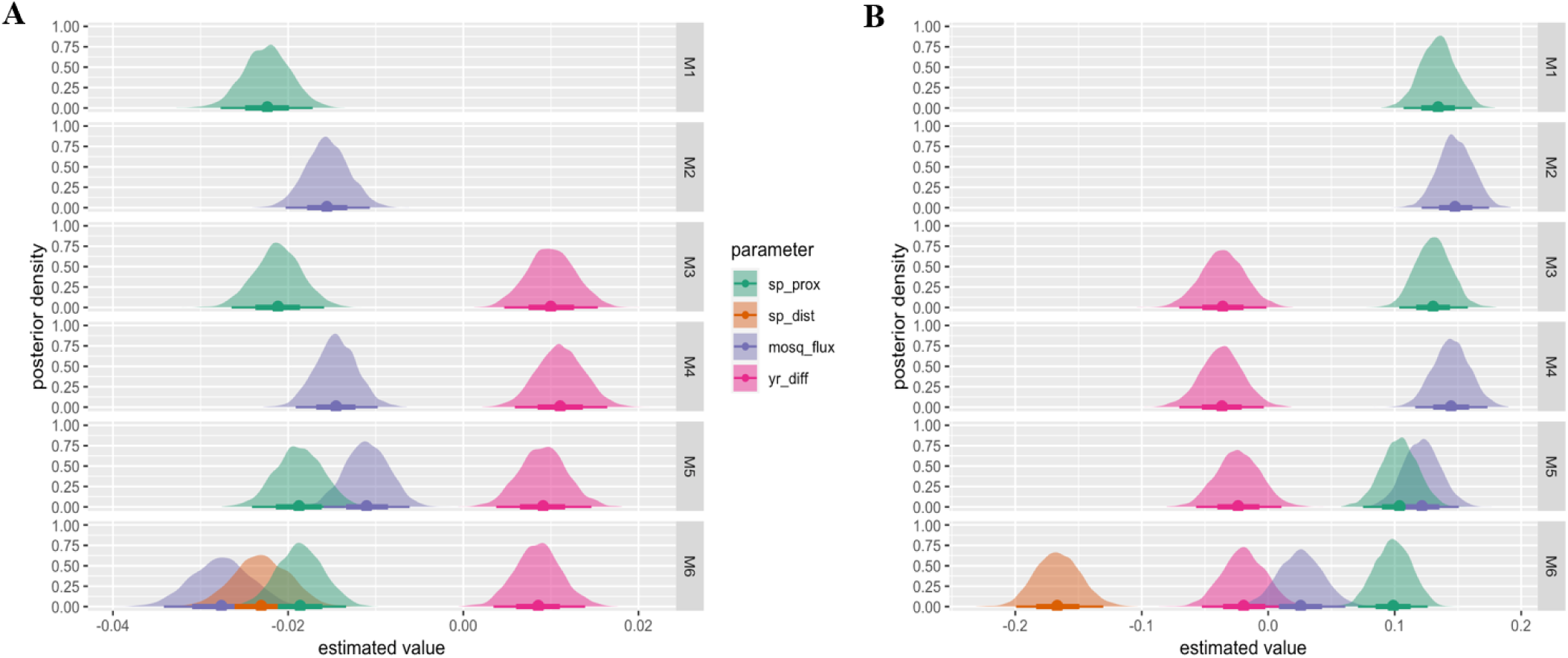
Estimated relationship between ITS2 pairwise genetic distance, spatial distance (sp_dist; geodesic distance between sample locations in meters), spatial proximity (sp_prox; measured as the negative exponential of distance), potential tiger mosquito flux (mosq_flux; estimated from commuting patterns and tiger mosquito population distribution) and temporal distance (yr_diff; measured as absolute difference between years in which samples were taken) on the beta mean (*μ*) parameter (A) and on the zeros (B) in the zero-inflated beta regression models. Parameters are estimated from a set of Bayesian multilevel zero-inflated Beta regressions with multiple-membership random intercepts for the samples and sampling years represented in each pair.

**Fig 6.**
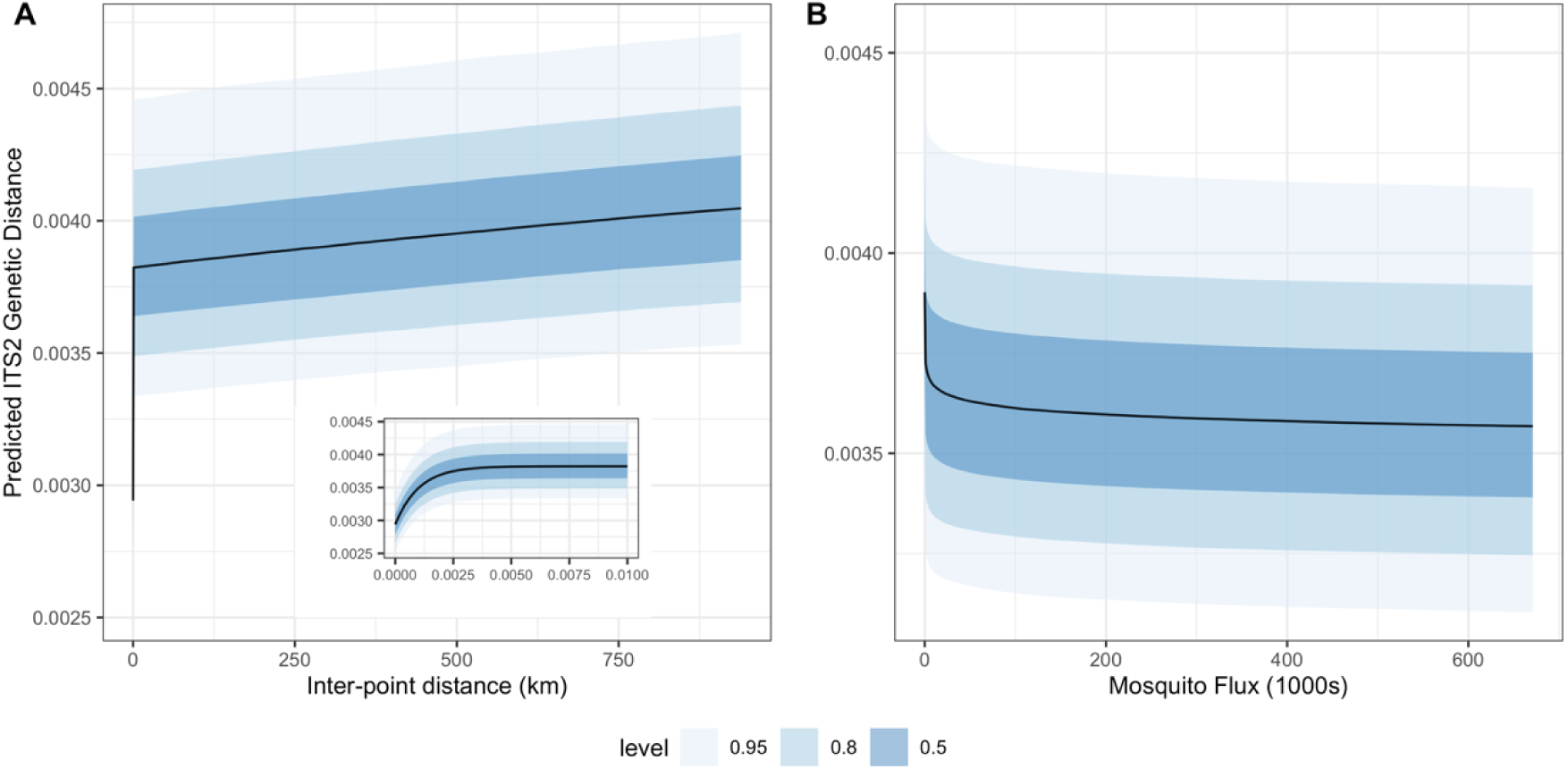
Predicted ITS2 pairwise genetic distance as a function of the spatial distance and spatial proximity variables taken together (A) and potential tiger mosquito flux (B) in the zero-inflated beta regression Model 6. Panel A shows predictions for the range of inter-point distances in the modelled data (0-940 km), holding potential mosquito flux at its observed median, setting the sampling years to 2011 and 2015, and arbitrarily selecting a sample pair and its associated provinces for purposes of the model’s random intercepts. The inset plot in this panel shows a close-up of the predictions at very small distances (0-10 m). Panel B shows predictions for the range of mosquito fluxes in the modelled data (0-672 km), holding inter-point distance at its median, setting the sampling years to 2011 and 2015, and arbitrarily selecting a sample pair and its associated provinces.

Fig 6A shows the effects predicted by M6 of changes in inter-point distance (reflected simultaneously in the spatial distance and spatial proximity variables) on ITS2 genetic distance. The range of inter-point distance values used for these predictions is the same as that observed in the data. Potential mosquito flux is held at its median, the sampling years are set to 2011 and 2015 (to give the widest range observed) and the sample pair (for purposes of the random intercepts) is arbitrarily selected. The overall pattern is non-linear, reflecting the combination of the spatial proximity and spatial distance variables. There is an initial steep increase in predicted ITS2 genetic distance for samples taken within a few meters of one another, which can be seen most clearly in the inset plot of Fig 6A. Beyond these highly proximate samples, predicted ITS2 genetic distance then rises more gradually, driven by the spatial distance variable.

Fig 6B shows the effects predicted by M6 of changes in potential mosquito flux on ITS2 genetic distance. The range of fluxes used for these predictions is the same as that observed in the data. Inter-point distance is held at its median and used for calculating the spatial proximity and spatial distance variables, the sampling years are set to 2011 and 2015 (to give the widest range observed) and the sample pair (for purposes of the random intercepts) is arbitrarily selected. Here we see a steep drop in predicted ITS2 genetic distance as potential mosquito flux increases from 0 to several hundred per day, with effect of increased mosquito fluxes becoming weaker at higher values, reflecting the log-linear relationship used in the model. Although in this case the pattern is overwhelmed by the uncertainty of the predictions (the wide posterior predictive distributions shown in the lighter shades of blue), this is the result of the combined uncertainty of the other variables in the model; the effect of potential mosquito flux, net of these other variables, is very clearly shown in the parameter estimates in Fig 5.

Fig 7 shows the combined effects of inter-point distance and potential tiger mosquito flux changing together. We see, first, how the lowest values of inter-point distance (bottom edge of plot) correspond with the lowest predicted ITS2 genetic distances while the lowest values of potential tiger mosquito flux (left edge of plot) correspond with the highest. When both variables are at their lowest (bottom left corner), inter-point distance is determinative: predicted ITS2 genetic distance is low here regardless of potential tiger mosquito flux. Beyond these lowest values, we see how predicted tiger mosquito flux acts to hold predicted ITS2 genetic distance down even as distance increases. Although predicted ITS2 genetic distances never reach their lowest values at these longer distances, the potential tiger mosquito fluxes of around 26,000 mosquitoes per day (this is the potential flux, for example, between Barcelona municipality and El Prat de Llobregat) keep the predicted ITS2 genetic distance within the range of 0.0035-0.0036, even at distances of 100 km.

**Fig 7.**
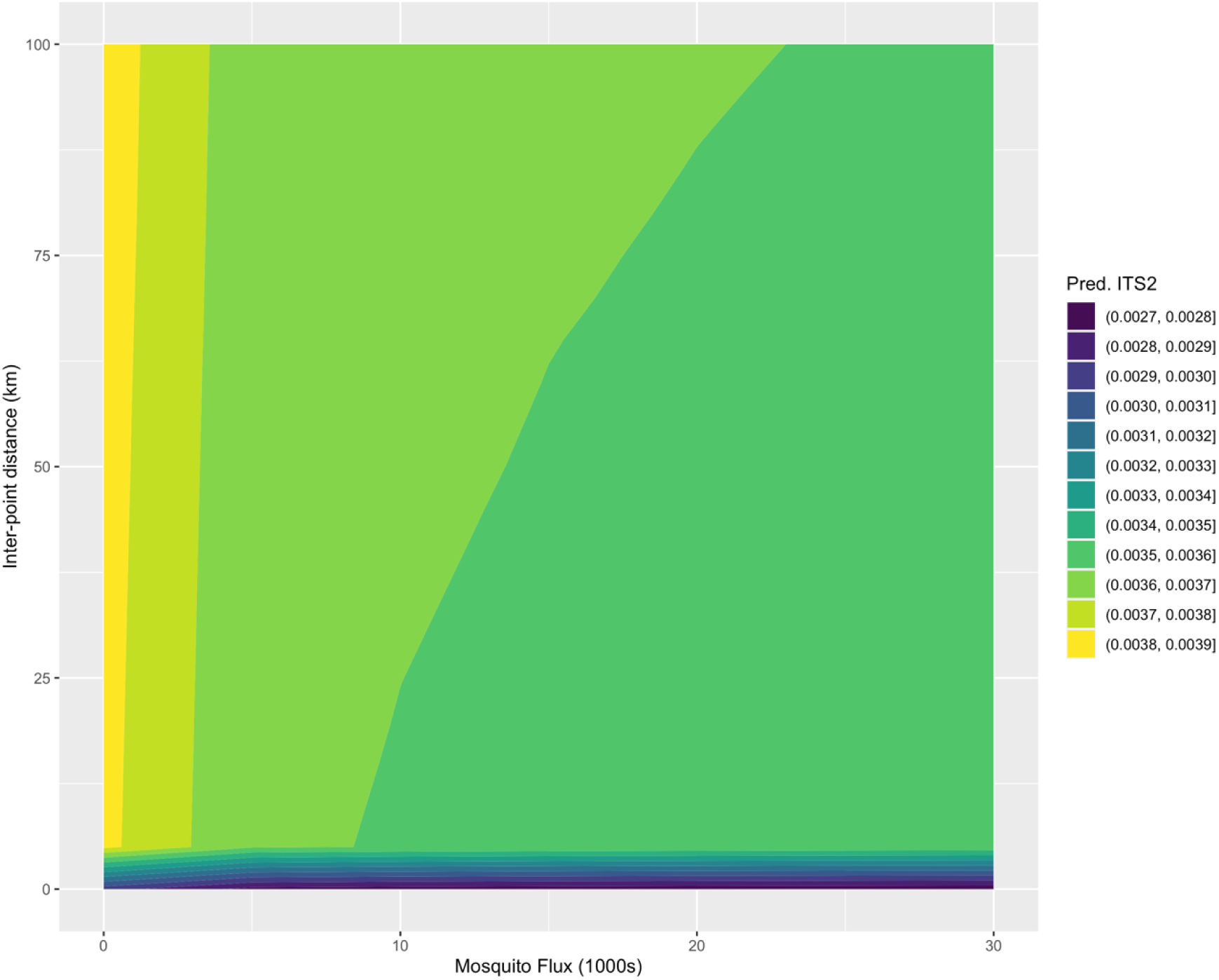
Predicted ITS2 pairwise genetic distance (indicated by fill colour) as a function of inter-point distance (the spatial distance and spatial proximity variables taken together) and potential tiger mosquito flux in the zero-inflated beta regression Model 6. Predictions are shown inter-point distances between 0 and 100 km and potential tiger mosquito fluxes between 0 and 30,000, setting the sampling years to 2011 and 2015, and arbitrarily selecting a sample pair and its associated provinces for purposes of the model’s random intercepts.

Note, finally, that although the ITS2 genetic distances and effect sizes shown in Figs 5-7 are small overall, this should be interpreted in light of the distribution of observed genetics distances, which ranged from 0 to only 0.07, with a standard deviation of 0.01.

## Discussion

*Aedes albopictus* has spread worldwide and particularly in Europe at a fast pace, making it one of the 100 most invasive species on Earth [12]. In Spain, after *Ae. albopictus* was first found near Barcelona in 2004 [18], a continuous spread along the Mediterranean coast was observed [17]. All Mediterranean provinces are currently colonised, along with the Basque Country in the northern coast and several inland territories [20], where the species continues to expand. In light of this, understanding *Ae. albopictus* dispersal routes across scales is crucial for planning effective early warning surveillance in non-invaded areas and implementing surveillance and control activities in the areas already colonised. The same information can also be valuable in predicting the transmission risk of pathogens by this vector. Knowledge of the effect of vehicles and transport infrastructures in the genetic structure of vector populations and their overall spreading capacity can comprehensively show our potential as a natural selective force and the existing contradictions between globalization and our efforts to combat biological invasions and pests [78].

Our models indicate that at very small spatial scales (i.e. several meters) the genetic variation measured by ITS2 is sharply reduced, likely representing seasonal mosquito pools that come from a main source. Beyond these scales, genetic variability steadily increases with spatial distance, as a clear positive correlation between dispersal distance and genetic variation appears. This is reflected in the combined effects of the (small scale) spatial proximity variable and the linear spatial distance variable. More interestingly, our models suggest that human transportation has a role in shaping *Ae. albopictus* nuclear genetic structure by means of passive dispersal of adult tiger mosquitoes in cars: there is a clear negative relationship between mosquito flux and genetic variation. Previous studies have also highlighted that, although *Ae. albopictus* has low natural dispersal capabilities, human-aided transport (especially cars) has probably facilitated significantly the tiger mosquito’s movement and invasion process [29, 30]. Our findings are consistent with these but go a step further by suggesting that the “hitchhiking” of *Ae. albopictus* in cars observed in Eritja et al. [30] actually helps to explain observed patterns of population genetics. While genetic variability increases with spatial distance, car transport can strongly reduce this effect. The high-resolution of our dataset makes it possible to show the significant role of human transportation in shaping the genetic constitution of *Ae. albopictus* and promoting regional gene flow. Mosquito movement can be affected by human activities like commuting and human-made structures like roads, which combined act as bridges for dispersal by favouring gene flow and promoting genetic mixing.

At a much broader spatial scale, we find a general lack of genetic structure and geographic consistency among haplotypes across the large expanse of the Iberian Peninsula, with numerous haplotypes shared among several distant areas (e.g. the most abundant ITS2 haplotypes found in Spain were also detected in several other European and Asian countries). This suggests that human-mediated large-scale dispersal of *Ae. albopictus* is also common, and point to a pattern of regular introductions of the species from abroad, through e.g. transportation of used tires and aquatic plants, which allows for survival and establishment at long distances of whole batches of eggs. Previous genetic studies of *Ae. albopictus* have also pointed out an apparent lack of genetic structure according to geography, both in the native and introduced range of the species [reviewed in 38]. Such a genetic feature is likely the consequence of the high level of human-mediated spread from several genetically distinct source populations followed by global dispersal, and it is concordant to what has been found in other wide-ranging invasive insect species, especially those that are closely associated with humans, e.g. the German and American cockroaches (*Blattella germanica* and *Periplaneta americana*, respectively) [79, 80] and the longhorn crazy ant *Paratrechina longicornis* [81]. Taken together, our results highlight the role of human activities in promoting unintentional mid-and long-distance dispersal and thus in shaping the current genetic structure of insect species commonly found in human-modified landscapes.

Our study has some limitations that should be considered. Firstly, the modelled genetic diversity range, which was obtained from ITS2 pairwise genetic distances, is low. This likely has to do with the relatively low variability of the analysed genetic marker. Indeed, nuclear genes, as well as mitochondrial fragments, are expected to be less variable and bear lower resolving power than highly mutating markers, e.g. microsatellites. Secondly, intragenomic heterogeneity (i.e. presence of multiple haplotypes within the same individual) of ITS2 has been reported in several mosquito species including the genus *Aedes* [82, 83], and this can pose a challenge in DNA sequencing and analysis. In this study, all individuals presenting intragenomic heterogeneity were thus excluded from analysis. As for future directions, although ITS2 has proved to be a useful marker for studies on the spread of *Ae. albopictus* [84], the reassessment of the species’ genetic diversity and population structure through the use of molecular markers with greater variability and/or potential, such as microsatellites and single nucleotide polymorphisms (SNPs), could provide more detailed and deeper insights into the description of fine-scale dispersal patterns, gene flow and introduction routes.

Global patterns of mtDNA and nuclear variation were highly discordant, with mtDNA showing little genetic diversity and a single star-like haplotype network. This is in agreement with previous studies showing *Ae. albopictus* levels of nuclear variation within the range of most insects, but extremely low mtDNA variation both within and among populations [85–88]. Interestingly, we found a high infection rate (79%) of *Wolbachia* in the studied *Ae. albopictus* samples. *Wolbachia* is a genus of maternally-inherited endosymbiotic bacteria that is known to induce male killing, feminization, parthenogenesis and cytoplasmic incompatibility, which facilitate its spread within the arthropod population [89]. *Wolbachia* is capable of inducing selective sweeps in mtDNA,

i.e. fixation of a single or few mtDNA haplotypes that may become widespread in the host population through cytoplasmic hitchhiking driven by *Wolbachia* invasions [35]. Selective sweeps on mtDNA have been shown to not only reduce haplotype diversity producing a characteristic single star-like network, but also to cause the remaining set of haplotypes to deviate from neutrality [34]. Within Culicidae, the natural presence of *Wolbachia* has been documented in more than 30 species [e.g. 50, 51, 90, 91], with *Ae. albopictus* harbouring significantly lower mtDNA diversity than the uninfected species [90]. In light of this, our results point to *Wolbachia* as causative agent for the lack of mitochondrial polymorphism here recovered, as suggested for other insect species, e.g. *Acraea* butterflies [92] and the cherry fruit fly *Rhagoletis cerasi* [93]. Nevertheless, the low variability in mtDNA in the introduced ranges of *Ae. albopictus* could also be caused by demographic processes, such as genetic drift or population bottleneck during rapid colonization [94]. However, demographic processes cause changes in variation for both mitochondrial and nuclear markers, even though mtDNA is expected to show a stronger response [95]. Furthermore, the lack of mtDNA variation was also found in the native range of *Ae. albopictus* [85, but see 96], strengthening our hypothesis of a *Wolbachia*- induced selective sweep. Alternatively, we cannot rule out that the low COI diversity here detected may be at least partially due to the length of the analysed fragment, as higher diversity has been observed when targeting larger COI fragments (>1300 bp) [97, 98]. Further research is needed to test this hypothesis.

Mitochondrial DNA has been extensively used to shed light on the geographic origin of invasive *Ae. albopictus* populations and arthropods in general [86, 87, 99], and disentangle their phylogeographic history [96, 100]. Nevertheless, there is broad recognition that mtDNA can commonly be under selection, thus challenging the assumption of neutrality postulated by several population, phylogeographic and phylogenetic studies [32, 101, 102]. Selection can arise due to e.g. mito-nuclear co-evolution, adaptation of mtDNA to different climatic/environmental conditions, and selective sweeps of beneficial mtDNA haplotypes [102]. Mitochondrial DNA is a single, linked molecule with low or no recombination, meaning that selective processes such as selective sweeps can have a profound impact on the apparent rate of genetic drift [32]. Therefore, we suggest caution should be used in drawing conclusions from mtDNA alone and, whenever possible, aim for a multilocus approach to achieve a correct understanding of the genetic population structure and history of the study species.

## Acknowledgements

We would like to thank those citizen scientists participating in the *Mosquito Alert* project who captured and sent us adult tiger mosquitoes for this study. We are also thankful to Victor Ojeda, Sandra Serra and Marc Pradell for their valuable help at the initial stage of the study.

## Supporting information

**S1 Fig. Map of potential tiger mosquito flux between municipalities in Spain.** Brighter yellow represents higher flux. Lines drawn only between municipality pairs with estimated flux of at least two tiger mosquitoes per day.

**S2 Fig. Comparison of zero-inflated beta regression models based on Bayesian R- squared.** Points indicate the R-squared estimate for each model (M1-M6). Lines indicate the 95% credible interval.

**S3 Fig. Posterior distributions for year random intercepts in the zero-inflated beta regression models (M1-M6) of the beta mean (*μ*).**

**S4 Fig. Posterior distributions for year random intercepts in the zero-inflated beta regression models (M1-M6) of the zeros.**

**S1 Table. Comparison of zero-inflated beta regression models based on expected log pointwise predictive density (ELPD) using leave-one-out cross-validation (LOO).** Abbreviations: elpd_diff – ELPD point difference relative to Model 6 (M6); se_diff – Standard Error point difference relative to M6.

**S2 Table. Parameter estimates for the six zero-inflated beta regression models (M1-M6) and comparison of models based on Bayesian R-squared and expected log pointwise predictive density (ELPD) using leave-one-out cross-validation (LOO).** Abbreviations: Int. - random intercepts; sp_prox - spatial proximity;

ln_mosq_flux – natural log of potential tiger mosquito flux plus 1; yr_diff – temporal distance; sp_dist - spatial distance; SE - Standard Error.

